# The effect of plasma lipids and lipid lowering interventions on bone mineral density: a Mendelian randomization study

**DOI:** 10.1101/480426

**Authors:** Jie Zheng, Marie-Jo Brion, John P. Kemp, Nicole M. Warrington, Maria-Carolina Borges, Gibran Hemani, Tom Richardson, Zhen Qiao, Philip Haycock, Mika Ala-Korpela, George Davey Smith, Jon H. Tobias, David M. Evans

## Abstract

Statin treatment increases bone mineral density (BMD) and reduces fracture risk, but the underlying mechanism is unclear. We used Mendelian randomization (MR) to assess whether this relation is explained by a specific effect in response to statin use, or by a general effect of lipid-lowering. We utilized 400 single nucleotide polymorphisms (SNPs) robustly associated with plasma lipid levels and results from a heel BMD GWAS (derived from quantitative ultrasound) in 426,824 individuals from the UK Biobank. We performed univariate and multivariable MR analyses of low-density lipoprotein cholesterol (LDL-C), high density lipoprotein cholesterol (HDL-C) and triglyceride levels on BMD. To test whether the effect of statins on BMD was mediated by lowering lipid levels, MR was repeated with and without SNPs in the *HMGCR* region, the gene targeted by statins. Univariate MR analyses provided evidence for a causal effect of LDL-C on BMD (*β* = −0.060; −0.084 to −0.036; P = 4×10^-6^; standard deviation change in BMD per standard deviation change in LDL-C, with 95% CI), but not HDL or triglycerides. Multivariable MR analysis suggested that the effect of LDL-C on BMD was independent of HDL-C and triglycerides, and sensitivity analyses involving MR Egger and weighted median MR approaches suggested that the LDL-C results were robust to pleiotropy. MR analyses of LDL-C restricted to SNPs in the *HMGCR* region showed similar effects on BMD(*β* = −0.083; −0.132 to −0.034; P = 0.001) to those excluding these SNPs (*β* = −0.063; −0.090 to −0.036; P = 8×10^-6^). Bidirectional MR analyses provided some evidence for a causal effect of BMD on plasma LDL-C levels. Our results suggest that effects of statins on BMD are at least partly due to their LDL-C lowering effect. Further studies are required to examine the potential role of modifying plasma lipid levels in treating osteoporosis.

## Introduction

Statins, the 3-hydroxy-3-methylglutaryl coenzyme A (HMG-CoA) reductase inhibitors, are principal therapeutic agents in lowering blood cholesterol, especially, low density lipoprotein cholesterol (LDL-C). Several randomized controlled trials (RCTs) have reported increased bone mineral density (BMD) following statin administration (1)(2). These results could reflect a direct effect of statins on BMD, as suggested by findings from several *in vivo* studies that statins stimulate bone formation (3)(4)(5). An alternative possibility is that the relationship between statin use and BMD is at least partially mediated by an effect of LDL cholesterol on bone metabolism (6)(7). For example, Parhami et al proposed that products of lipid and lipoprotein oxidation may contribute to the pathophysiology of osteoporosis (8). Consistent with this hypothesis, several observational epidemiological studies have documented a link between coronary heart disease and osteoporosis (9)(10). However, observational epidemiological studies are subject to confounding and reverse causality, making interpretation of such associations difficult and their meaning uncertain.

Mendelian randomization (MR) uses genetic variants as instrumental variables to estimate the causal effect of modifiable environmental exposures on medically relevant outcomes (11)(12). For example, we recently used this method to demonstrate a causal effect of greater fat mass on BMD in children (13). In order to investigate the relationship between blood lipids and BMD, we performed a two sample MR study (14). We utilized summary GWAS data from the Global Lipids Genetics Consortium (15) as exposures, and summary GWAS data of ultrasound derived heel BMD on 426,824 UK Biobank European participants as the outcome (16). To obtain estimates of the causal effect of blood lipids on BMD, we performed two sample inverse variance weighted (IVW) MR (17), and also a series of sensitivity analyses including MR Egger regression (18), weighted median MR analysis (19) and multivariable MR (20)(21) which may produce more robust causal estimates in the presence of horizontal genetic pleiotropy (22). To determine whether any relationship between BMD and blood lipids might be mediated by direct effects of statin use, we compared causal estimates obtained using SNPs in the *HMGCR* gene (i.e. whose product is the target of statin therapy) versus estimates using SNPs outside this gene. Finally, we investigated whether there was any evidence for reverse causality (i.e. BMD causally influencing plasma lipids) by performing bidirectional MR (23).

## Methods

### Two sample inverse variance weighted Mendelian randomization analysis

We performed a series of two-sample MR analyses using summary results data from the Global Lipids Genetics Consortium (N = 331,368) (15) and the UK Biobank Study of BMD (N = 426,824) (16). In total, 400 conditionally independent SNPs robustly associated with blood lipids (P < 5 × 10^-8^) were selected as instruments for the MR analyses (see Supplementary Table 1). Of these SNPs, 195 variants were associated with HDL-C, 147 were associated with LDL-C, and 163 were associated with triglycerides at genome-wide levels of significance (see Supplementary Table 2, 3 and 4). We refer to MR analyses involving all these variants as analyses using the “Complete Set” of SNPs. To obtain estimates of the causal effect of lipid fractions on BMD, we performed two-sample IVW MR analysis (24) on each lipid fraction separately. In addition, adiposity has been reported to be causally related to BMD (13). To control for the possible introduction of confounding by including adiposity related variants in our MR analyses, we cross referenced our LDL-C instruments with the most up-to-date list of body mass index (BMI) related SNPs (25). We excluded LDL-C SNPs that were in linkage disequilibrium with known BMI variants (r^2^>0.5 in the 1000 Genome Europeans; Supplementary Table 3) and conducted the IVW MR analysis again using the remaining 137 instruments. Analyses were performed using the TwoSampleMR R package of MR-Base (26) (https://github.com/MRCIEU/TwoSampleMR).

### Sensitivity analyses

Standard MR analyses rely on the validity of a number of core assumptions to produce accurate estimates of the causal effect of the exposure on the outcome (11) (24) (27). One of these assumptions states that genetic instruments (lipid SNPs) must only potentially be related to the outcome (i.e. BMD) through their relationship with the exposure (lipid levels). Thus, if there are additional pleiotropic paths between the SNP and outcome that do not pass through lipid levels, then standard MR analyses may produce biased estimates of the causal effect of blood lipids on BMD. We therefore applied two recent extensions of the basic IVW MR method, that are more robust to the presence of genetic pleiotropy, MR Egger regression (18) and weighted median MR (19).

In MR Egger regression, estimates of the SNP-outcome association are regressed on estimates of the SNP-exposure association. As long as certain assumptions are met (see below), the slope of the weighted regression line provides an estimate of the causal effect of the exposure on the outcome that is free from the effects of horizontal pleiotropy (18). The intercept of the MR Egger regression line quantifies the amount of directional pleiotropy present in the data averaged across the different variants. Thus, the presence of overall directional pleiotropy can be assessed through statistical tests of the degree to which the intercept differs from zero. The validity of MR Egger regression critically rests on the ‘INSIDE assumption’ (INstrument Strength is Independent of Direct Effect) which states that across all SNPs there should be no correlation between the strength with which the instrument proxies the exposure of interest, and its degree of association with the outcome via pathways other than through the exposure (18). This is a weaker requirement than the no pleiotropy assumption in IVW MR analysis, so MR Egger regression is likely to be more robust to horizontal pleiotropy than standard IVW MR, although this comes at the cost of decreased power to detect a causal effect (18). In the context of the present study, provided the underlying assumptions are met, the slope of the MR Egger regression analysis should yield an estimate of the causal effect of LDL-C (or HDL-C or triglycerides) on BMD that is free from any confounding effects due to horizontal pleiotropy.

We assessed the no measurement error (NOME) assumption in MR Egger regression using an adaptation of the I^2^ statistic to the two-sample summary data MR context, which is referred to as I^2^_GX_ and accounts for uncertainty in the SNP-exposure estimates. I^2^_GX_ provides an estimate of the degree of regression dilution in the MR-Egger causal estimate (28).

We also used the weighted median MR approach as another sensitivity analysis, which provides consistent estimates of the causal effect of exposure on outcome even when up to 50% of the information contributing to the analysis comes from genetic variants that exhibit pleiotropy (19). Thus, MR Egger regression and weighted median MR provide causal estimates of the exposure (lipids) on the outcome (BMD) under different assumptions. If all approaches (i.e. IVW MR, MR Egger regression and weighted median MR) provide similar estimates of the causal effect of lipids on BMD, then we can be more confident that our findings are robust. All sensitivity analyses were performed using the MR-Base R package as described above (26).

When applying MR, we make an assumption that SNPs used to proxy lipids exert their primary association on lipids, and that any correlation with BMD is a consequence of a causal effect of lipids on BMD. However, if in reality BMD exerts a causal effect on lipids, then it is possible that some SNPs primarily associated with BMD might also reach genome-wide significance in a large GWAS of lipids. These BMD SNPs could then mistakenly be used as instruments for lipids, when in fact they should be used as instruments for BMD levels. In other words, in very large GWAS it can be difficult to determine whether a SNP has its primary association with the exposure under study, or the outcome. This was the case in the GLGC and UKBB studies as some of the BMD associated SNPs were also strongly associated with lipids (i.e. at genome-wide levels of significance). We therefore applied MR Steiger filtering (29) as implemented in the TwoSampleMR R package (30) to test the causal direction of each of the 400 lipid associated SNPs on the hypothesized exposures (lipids) and outcome (BMD). This approach infers the causal direction between phenotypes using a simple inequality. Given trait A causes trait B then one would expect that:

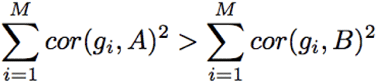

because cor(g_i_, B)^2^ = cor(A, B)^2^ * cor(g_i_, A)^2^, where “cor” denotes correlation, and the vector g contains a set of M SNPs that influence trait A. For any SNP that had a cor(g,A)^2^ < cor(g,B)^2^ (which means it showed evidence of primarily affecting BMD rather than lipids), we removed those lipids SNPs and conducted IVW MR, MR Egger and Weighted Median MR using the remaining instruments (“Steiger filtered” set). The process of choosing validated instruments using Steiger filtering followed these steps:

1. Select lipid instruments from the main analysis (p-value threshold 5×10^-8^).
2. Classify instruments in each MR analysis based on Steiger filtering:

- ‘TRUE’: evidence for causality in the expected direction i.e. lipids influence BMD.
- ‘FALSE’: evidence for causality in the reverse direction i.e. BMD influences lipids. Instruments with ‘FALSE’ were removed from the sensitivity analysis.
- ‘NA’: no result (due to missing effect allele frequencies in the outcome data or missing numbers of cases and controls for binary traits).

Individual Steiger filtering results can be found in Supplementary Table 2, 3 and 4.

### Two sample multivariable Mendelian randomization analysis

Since many of the SNPs used in the previous MR analyses were associated with more than one lipid fraction, we used multivariable MR (Figure 1) to estimate the causal effect of HDL-C, LDL-C and triglycerides on BMD, using a “weighted regression-based method” approach where the inverse-variance weights were applied to a multivariable regression model (20)(21). Multivariable MR has an advantage over univariate MR in that it accounts for the potential pleiotropic influence of other exposures included in the analysis (i.e. HDL-C, LDL-C and triglycerides). However, similar to IVW MR, multivariable MR makes a critical assumption that the relationship between the genetic instruments and the outcome is mediated exclusively by the exposure variables considered in the analysis (i.e. LDL-C, HDL-C and triglycerides), which of course may not be the case in reality. We therefore fitted a multivariable MR model with an unconstrained intercept term, which has the effect of allowing for directional pleiotropy, similar to the situation in MR Egger regression (31). Since multivariable MR does not require each genetic instrument to be related to every exposure variable (merely that each SNP is a strong instrument for at least one exposure), we applied the method to the complete set of 400 lipid associated SNPs. We performed sensitivity analyses coding the direction of the SNPs as LDL-C increasing, HDL-increasing and then triglceride-increasing to examine whether the direction of coding affected the multivariable MR Egger regression results.

**Figure 1.**
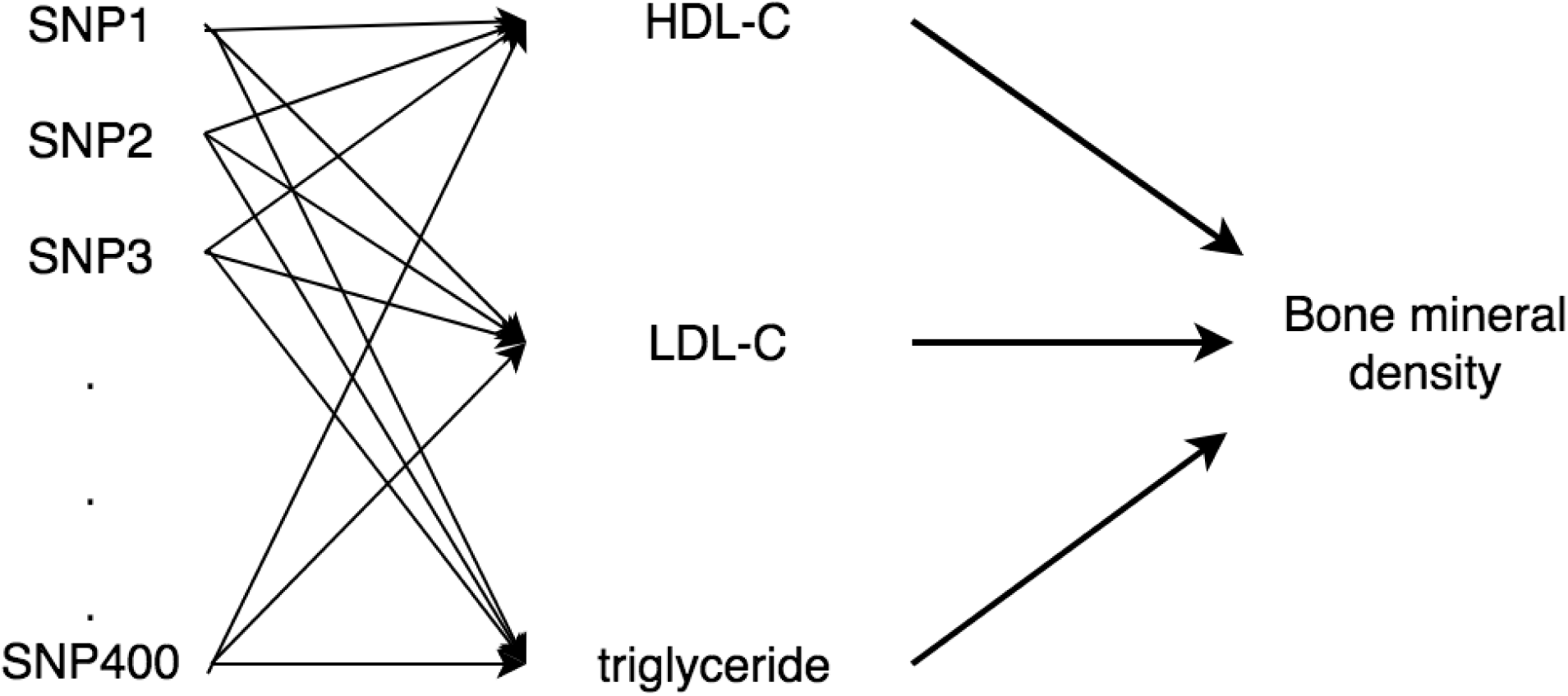
Directed acyclic graph illustrating multivariable MR analysis. In total, 400 conditionally independent SNPs robustly associated with blood lipids (p < 5 × 10^-8^) were used in the multivariable MR analysis. Many of the SNPs act pleiotropically and affect more than one lipid fraction. In addition, estimating an intercept in the multivariable MR regression (rather than having it constrained to zero) is akin to allowing for the possibility of additional horizontal pleiotropy in MR Egger regression.

### Predicting the impact of lipid lowering pharmaceutical interventions on bone mineral density

Analogous to what has been done in several previous MR studies of lipids and coronary heart disease (32)(33)(34), we used a selection of genetic variants at the 3-Hydroxy-3-Methylglutaryl-CoA Reductase *(HMGCR)*, Niemann-Pick C1-Like 1 (*NPC1L1*) and Proprotein convertase subtilisin/kexin type 9 (*PCSK9*) genes to mimic the expected action of statins, ezetimibe and evolocumab, respectively on BMD. Since some of the SNPs within these genes were in incomplete linkage disequilibrium (LD), we used a likelihood-based two-sample MR approach that takes into account the correlation between genetic instruments when estimating the causal effect of lipid lowering drugs on BMD (35). LD correlation estimates (r) between markers were obtained in CEU individuals using the LD matrix webserver (36). As a further sensitivity analysis, we repeated MR analyses using all SNPs outside the *HMGCR* region.

If statins causally affect bone mineral density via “direct effect” (i.e. independent of lipids), then we would expect to see significant causal estimates for MR analyses involving *HMGCR* SNPs, but not for analyses involving SNPs in the *NPC1L1* and *PCSK9* genes, nor the rest of the genome. In contrast, if the effect of the SNPs on BMD were mediated through blood lipids, then we would expect to obtain significant causal estimates using lipid associated SNPs across the rest of the genome. We formally compared the different causal estimates obtained using SNPs in the different gene regions using heterogeneity tests (37).

### Bidirectional Mendelian randomization

Finally, in order to investigate the possibility that BMD causally affects levels of blood lipids, we used summary results data from 1,410 conditionally independent autosomal variants reported in a recent BMD GWAS using 426,824 UK Biobank European individuals (16) as instrumental variables. We looked up these variants in summary results GWAS data on LDL-C, HDL-C and triglycerides from the Global Lipids Genetics Consortium (15)(34). We performed IVW MR, weighted median MR and MR-Egger regression methods using the TwoSampleMR R package as described above (26).

When applying Bidirectional MR, we also applied the Steiger filtering analysis (29) to test the causal direction of each BMD associated SNP on the hypothesized exposure (BMD) and outcomes (lipids). When it showed evidence of primarily affecting lipids rather than BMD, we removed those BMD SNPs and conducted bidirectional MR using the remaining instruments.

## Results

### Mendelian randomization estimates the causal effects of plasma lipids on BMD

Table 1 presents results from the univariate MR analyses of plasma lipids and BMD. Each regression coefficient represents the estimated causal change in standard deviations (SD) of BMD per SD change in serum level of HDL-C, LDL-C or triglycerides. IVW MR, MR Egger and weight median MR using the complete set of SNPs all suggested a causal effect of increased LDL-C on reduced BMD (IVW estimate: *β* = −0.060, 95%CI = −0.084 to −0.036, P = 4×10^-6^). Excluding SNPs related to BMI provided further evidence of a causal effect of increased LDL-C on reduced BMD (*β* = −0.058, 95%CI = −0.082 to −0.034, P= 3×10^-6^) (Supplementary Figure 1). In contrast, univariate MR analyses revealed little evidence for a causal effect of HDL-C or triglycerides on BMD (IVW HDL-C estimate: *β* = −0.016, 95%CI = −0.40 to 0.08, P = 0.2; IVW triglycerides estimate: *β* = 0.021, 95%CI = −0.49 to 0.07, P = 0.1). The Steiger filtering analysis suggested that most of the lipids SNPs exerted their primary effect on lipids as opposed to BMD. 146 SNPs showed evidence of a primary causal effect on LDL-C, 190 SNPs on HDL-C and 158 SNPs on triglycerides opposed to BMD (Table 1). MR using the Steiger filtered set of SNPs also showed strong evidence of LDL-C causally influencing BMD (IVW estimate: *β* = −0.058, 95%CI = −0.081 to −0.035, P = 8.9×10^-7^) and there was little evidence that HDL-C or triglyceride levels may influence BMD (IVW HDL-C estimate: *β* = −0.013, 95%CI = −0.31 to 0.05, P = 0.15; IVW triglycerides estimate, *β* = 0.015, 95%CI = −0.035 to 0.05, P = 0.13) (Table 1). Funnel plots and scatter plots for this sensitivity analysis are presented in Supplementary Figure 2.

**Table 1.** Summary of the univariate Mendelian randomization estimates of the causal effect of plasma lipids on BMD using different sets of SNPs as instruments. BMD was the outcome for all analyses.

| Exposure | SNP selection | Methods | N_SNPs | Beta | Standard error | P value |
| --- | --- | --- | --- | --- | --- | --- |
| LDL-C | complete | IVW | 147 | -0.06 | 0.012 | $4 \times 10^{-6}$ |
| | complete | WM | 147 | -0.028 | 0.007 | $1 \times 10^{-4}$ |
|  | complete | Egger | 147 | -0.04 | 0.018 | 0.03 |
| | Steiger filtered | IVW | 146 | -0.058 | 0.012 | $8.9 \times 10^{-7}$ |
| | Steiger filtered | WM | 146 | -0.027 | 0.007 | $1.3 \times 10^{-4}$ |
|  | Steiger filtered | Egger | 146 | -0.043 | 0.017 | 0.01 |
| HDL-C | complete | IVW | 195 | -0.016 | 0.012 | 0.20 |
|  | complete | WM | 195 | 0.009 | 0.005 | 0.06 |
|  | complete | Egger | 195 | 0.023 | 0.016 | 0.10 |
|  | Steiger filtered | IVW | 190 | -0.013 | 0.009 | 0.15 |
|  | Steiger filtered | WM | 190 | 0.010 | 0.005 | 0.03 |
|  | Steiger filtered | Egger | 190 | 0.019 | 0.012 | 0.11 |
| Triglycerides | complete | IVW | 163 | 0.021 | 0.014 | 0.10 |
|  | complete | WM | 163 | 0.007 | 0.007 | 0.30 |
|  | complete | Egger | 163 | -0.004 | 0.02 | 0.80 |
|  | Steiger filtered | IVW | 158 | 0.015 | 0.010 | 0.13 |
|  | Steiger filtered | WM | 158 | 0.006 | 0.006 | 0.31 |
|  | Steiger filtered | Egger | 158 | 0.006 | 0.014 | 0.69 |

Abbreviations: ‘complete’ refers to all SNPs associated with the particular lipid fraction, “Steiger filtered” refers to SNPs associated with the lipid fraction and pass Steiger filtering (29); Beta represents the standard deviation change in BMD per standard deviation change in plasma lipid level; N\_SNPs refers to the number of SNPs used as instrumental variables in the analysis; HDL-C: high density lipoprotein cholesterol; LDL-C: low density lipoprotein cholesterol; BMD: bone mineral density; IVW: inverse variance weighted Mendelian randomization; Egger: MR-Egger regression; WM: weighted median Mendelian randomization.

The funnel plots presented in Figure 2 display MAF-corrected genetic associations for each of the individual SNPs on lipid levels (y-axis) plotted against their causal effect estimates (x-axis). Visual inspection of Figure 2 provided little indication for the existence of directional pleiotropy for LDL-C (panel A), but a suggestion of directional pleiotropy for HDL-C (panel B) and potentially for triglycerides (panel C). In particular, SNPs less strongly related to increased HDL-C tended to be associated with reduced BMD. This interpretation was consistent with estimates of the intercepts from the MR Egger regression analyses (LDL-C: intercept = −0.001, P = 0.1; HDL-C: intercept = −0.004, P = 3×10^-4^; Triglycerides: intercept = 0.002, P = 0.08). Figure 2 also illustrates the associations between the LDL-C (panel D) / HDL-C (panel E) / triglycerides (panel F) variants and BMD in the form of scatter diagrams, with the MR Egger regression and IVW MR lines superimposed on the data points (the slopes representing the estimated causal effects).

**Figure 2.**
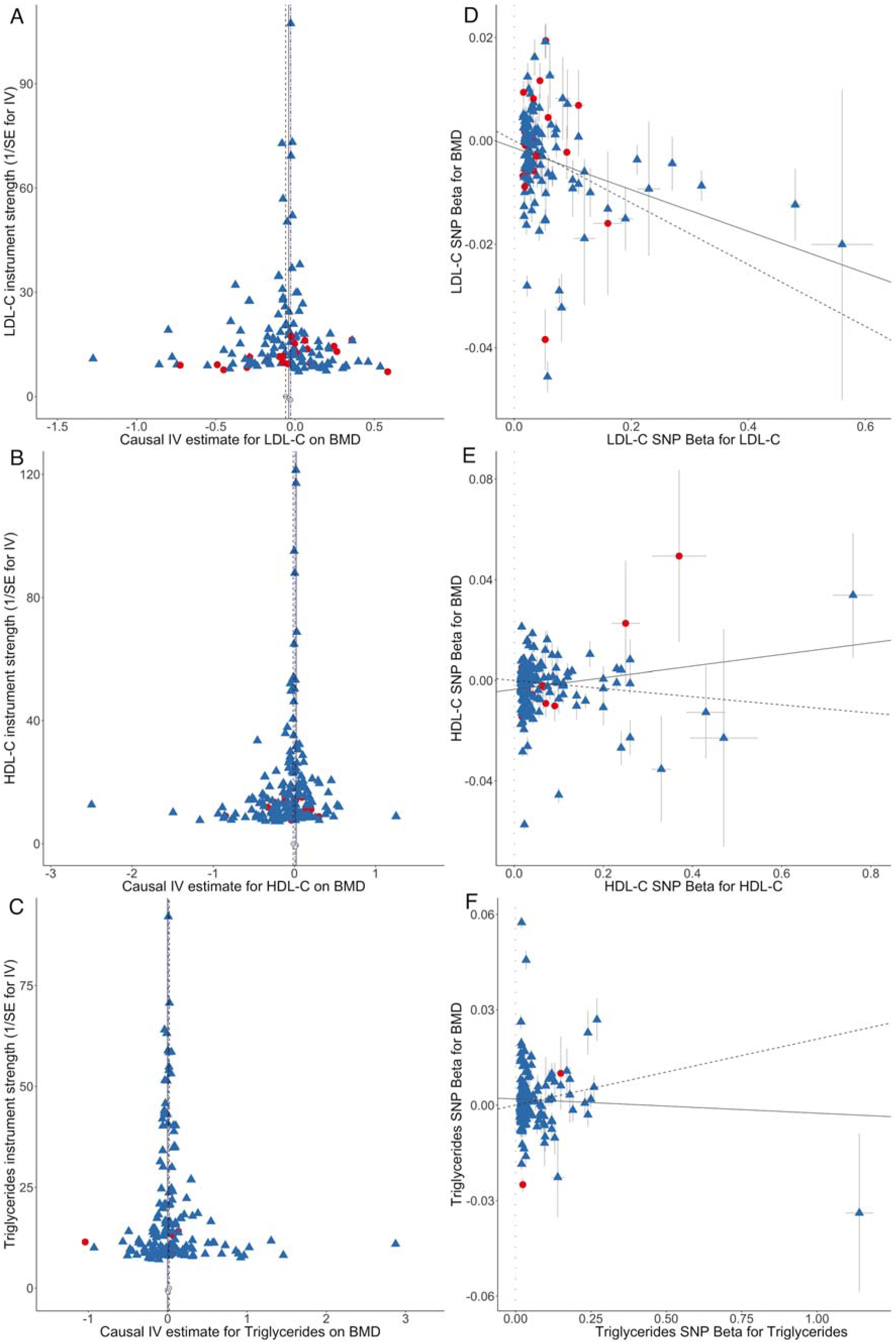
Results of the MR analysis of lipid levels on BMD using the complete set of instruments. Funnel plots displaying instrument strength (y-axis) plotted against causal effect estimates (x-axis) for SNPs associated with LDL-C (Panel A), HDL-C (Panel B), and triglycerides (Panel C) and scatter plots displaying estimates of the association between each SNP and BMD (y-axis) against estimates of the association between each SNP and lipid level, i. e. LDL-C (panel D), HDL-C (panel E) and triglycerides (panel F). The error bars on each of the points represent, 95% CI. SNPs in red circles denote those associated with one lipid subtype (P<5×10^-8^) but not the other two (P>0.05), whereas the remaining SNPs are denoted by blue triangles. For Panel A, B and C, the inverse-variance weighted MR, MR-Egger and weighted median MR causal effect estimates are represented by dotted, solid and double dotted lines respectively. For Panel D, E and F, the slope of the solid line represents the MR-Egger regression estimate of the causal effect of serum lipids on BMD and the inverse-variance weighted estimate is represented by the slope of the dotted line. The y-intercept of the solid regression line is an estimate of the degree of directional pleiotropy in the dataset.

Assessment of the NO Measurement Error (NOME) assumption (28) with respect to the MR-Egger estimate gave unweighted I^2^_GX_ = 0.995 and weighted I^2^_GX_ = 0.995. This suggests a minor 0.5% attenuation of the causal estimate toward zero, as a consequence of uncertainty in the SNP exposure estimates.

Multivariable IVW analysis provided additional evidence for a causal effect of LDL-C on BMD (*β* = −0.055, 95%CI = −0.080 to −0.030, P = 2.7×10^-5^), independent of the effects of HDL-C and triglycerides. The causal effect estimate was comparable to estimates produced from the previous univariate IVW MR analyses (Figure 3). As shown in Supplementary Table 5, there was no independent association between HDL-C and BMD (*β* = −0.020, 95%CI = −0.046 to −0.006, P = 0.123) and triglycerides on BMD (*β* = 0.013, 95%CI = −0.028 to 0.043, P = 0.397), consistent with univariate MR findings (Figure 3). The direction the alleles were coded in the multivariable analyses did not materially affect the results (Supplementary Table 5).

**Figure 3.**
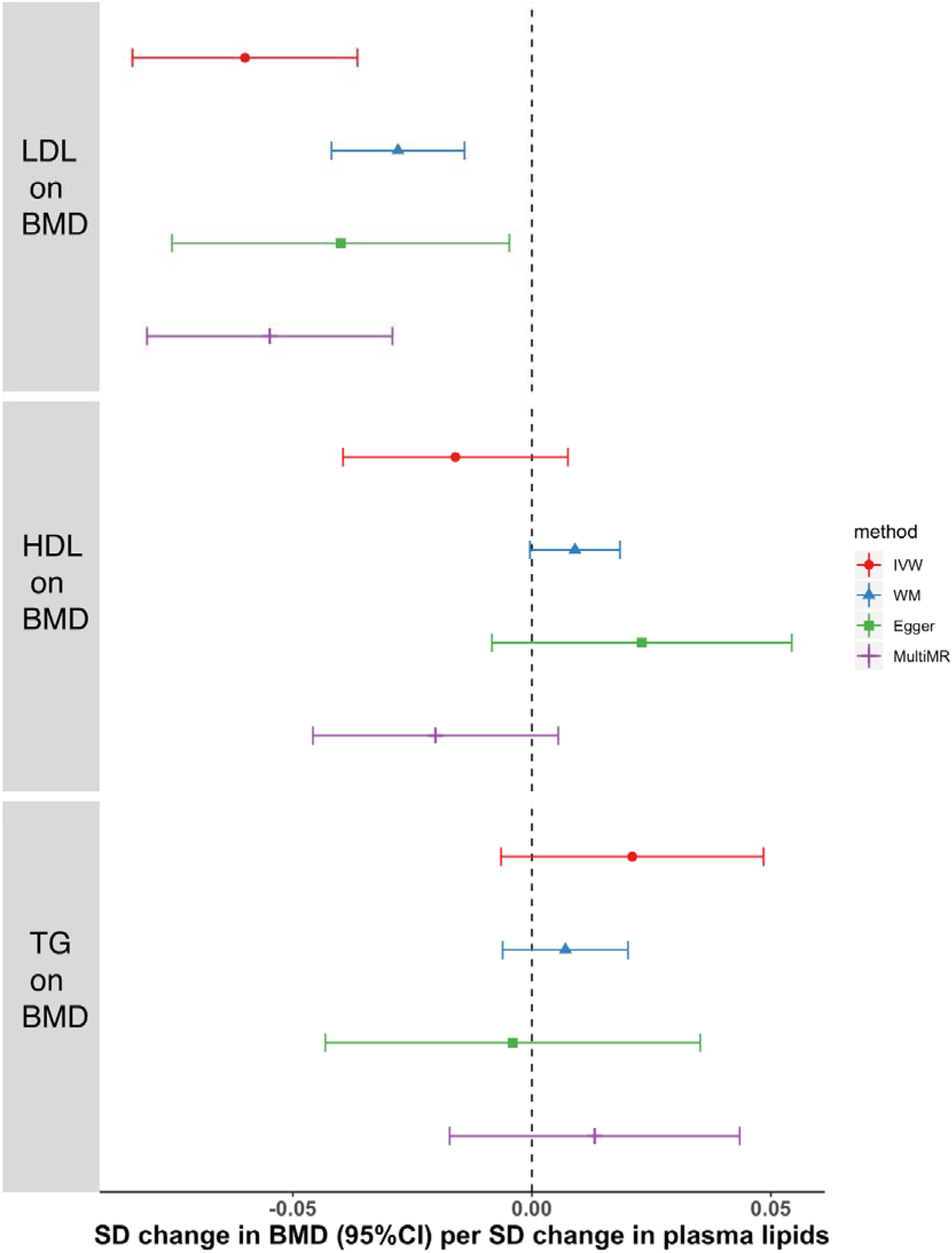
Forest plot comparing causal effect estimates of serum lipid levels on BMD using univariate and multivariable MR. The analysis was conducted using the complete set of lipid associated SNPs Abbreviations: SD, standard deviation; LDL-C: low density lipoprotein cholesterol; HDL-C: high density lipoprotein cholesterol; TG: triglycerides; BMD: bone mineral density; IVW inverse variance weighted; Egger, MR-Egger regression; WM, weighted median Mendelian randomization; MultiMR, multivariable Mendelian randomization. Note: all outlier exclusion approaches lead to reduced standard errors.

### Genetic prediction of the impact of lipid lowering interventions on bone mineral density

Table 2 displays estimates of the causal effect of LDL-C level on BMD obtained using SNPs from genes whose proteins are targets for lipid lowering drugs. Results obtained using 5 SNPs in the region of the *HMGCR* gene (33) suggest that reducing the activity of *HMGCR* (i.e. mimicking the effect of statins) increases BMD (*β* = −0.083, 95%CI = −0.132 to −0.034, P = 0.001). In contrast, results using 7 SNPs in the region of the *PCSK9* gene (33) plus rs11591147 (34), and 5 SNPs in the region of *NPC1L1* from Ference et al (32), suggested that genetically reducing the activity of *PCSK9* (mimicking Evolocumab) and *NPC1L1* (mimicking Ezetimibe) had no clear effect on BMD *(PCSK9: β* = −0.007, 95%CI = −0.027 to 0.013, P = 0.486; *NPC1L1: β* = −0.004, 95%CI = −0.059 to 0.051, P = 0.887). Interestingly, there was some evidence of heterogeneity in causal effect estimates across the different SNPs in the *PCSK9* gene, with one SNP in particular providing evidence for a causal effect in the opposite direction to the majority of the other SNPs. The SNPs used to explore the effect of the lipid lowering drugs are listed in Supplementary Table 6.

**Table 2.**
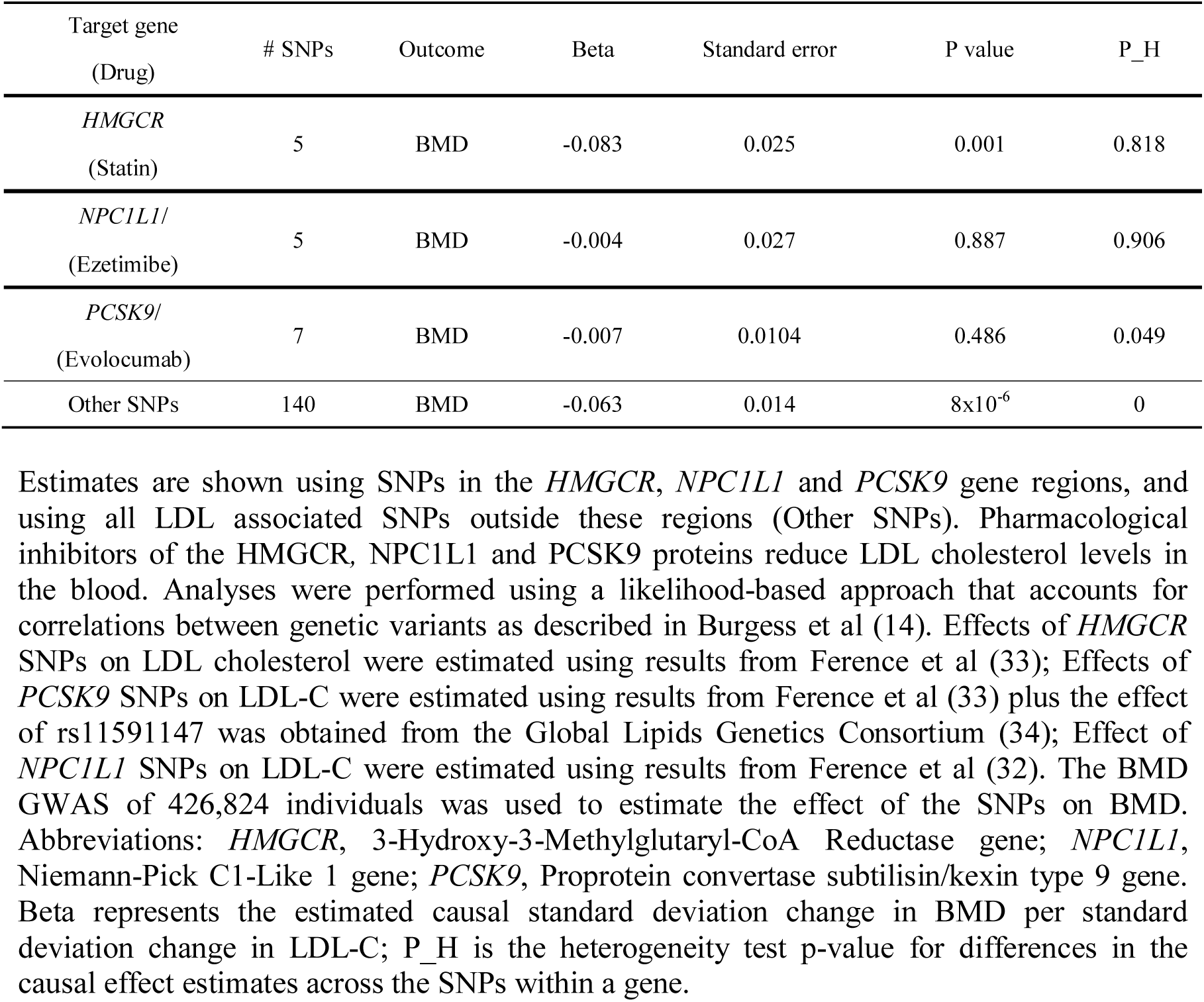
Instrumental variable estimates of the causal effect of LDL-C levels on BMD.

### Sensitivity analysis using non-HMGCR lipid lowering SNPs on bone mineral density

Table 2 and Supplementary Figure 3 display the causal estimates of LDL-C on BMD excluding the *HMGCR* SNPs from the MR analyses. We found that the 140 LDL-C associated SNPs outside the *HMGCR, PCKS9* and *NPC1L1* regions still produced significant estimates of a negative causal effect of LDL-C on BMD *β* = −0.063, 95%CI = −0.090 to - 0.036, P = 8×10^-6^). Supplementary Table 7 shows heterogeneity test results comparing causal estimates obtained from different gene regions (LD correlation matrices between SNPs in *HMGCR, PCSK9* and *NPC1L1* are shown in Supplementary Table 8). We found that the causal effect estimates using SNPs in the *HMGCR* gene were not different from those generated from the rest of the genome (excluding *HMGCR, PCSK9* and *NPC1L1* SNPs) (Cochran Q = 0.487, P=0.485). This finding suggests that some of the effect of the SNPs on BMD may be mediated through LDL-C (i.e. through mechanisms not involving HMGCR and statins). In addition, the confidence intervals surrounding the causal effect estimates obtained using SNPs in *PCSK9* and *NPC1L1* were wide and overlapped zero and were also different to the ones obtained using the *HMGCR* SNPs and the rest of the genome.

### Bi-directional Mendelian randomization estimates of the causal effect of BMD on plasma lipids

Finally, we investigated whether there was any evidence for BMD having a causal effect on blood lipids. 404 out of 1,410 SNPs reported as robustly associated with BMD from the UKBB study could be found in the blood lipids GWAS data (34). The Steiger filtering analysis suggested that most of these SNPs exerted their primary effect on BMD as opposed to lipid levels. 394 SNPs showed evidence of a primary causal effect on BMD as opposed to LDL-C, 389 SNPs opposed to HDL-C and 396 opposed to triglycerides (last four columns in Supplementary Table 9). Supplementary Table 10 presents univariate MR results for the effect of these remaining BMD associated SNPs on plasma lipids. IVW MR, weighted Median MR and MR-Egger regression results showed no strong evidence of BMD causally influencing HDL-C or triglyceride levels although there was some evidence that BMD might influence LDL-C. Interestingly, even after Steiger filtering and MR Egger regression, Cochrane Q statistics suggested the presence of considerable heterogeneity remaining in the analysis. The funnel plot and scatter plot for the bidirectional MR are presented in Supplementary Figure 4

## Discussion

We performed an MR study to examine whether previous findings of an inverse correlation between serum cholesterol and BMD reflected a causal relationship. We found that BMD increased by 0.064 SD per SD lower LDL-C, based on IVW MR analyses of the complete SNP set. Interestingly, MR analyses using 5 SNPs in the region of the *HMGCR* gene produced slightly higher estimates of the causal effect of LDL-C lowering on BMD. Taken together, these observations suggest that gains in BMD following statins are at least partly due to a causal effect of lowering LDL-C.

Aside from studies of statins (1)(2) (3)(4)(5), other evidence that LDL-C levels are inversely related to BMD is relatively lacking. For example, we are aware of only one small study examining the effect of the cholesterol-lowering agent ezetimibe on BMD, where no association was detected (38). Other cholesterol-lowering agents might lack the same tendency as statins to reduce BMD, which is further suggested by the results of our sensitivity analyses in which genetic instruments for other classes of cholesterol-lowering agents, ezetimibe and evolocumab, were unrelated to BMD. However, this finding contrasts with the effects of non-statin pathways as a whole on BMD, assessed by examining all LDL-C genetic instruments apart from *HMGCR*, which showed similar effects to instruments related to *HMGCR*. Conceivably, ezetimibe and evolocumab might exert distinct, adverse effects on BMD, countering beneficial effects resulting from lowering of LDL-C levels.

In terms of observational analyses, consistent with our results, a recent meta-analysis of 10 studies found that LDL-C levels were higher in patients with osteoporosis compared to controls (39). Such a relationship could conceivably contribute to the association between coronary vascular disease, for which high LDL-C levels are an important risk factor, and osteoporosis (9)(10). An inverse relationship between LDL-C and BMD, suggested by our results, is consistent with evidence that lipids may contribute to the pathophysiology of osteoporosis through lipid oxidation (7). For example, oxidized lipids, especially oxidized LDL, characteristic of hyperlipidemia, may have direct adverse effects on cellular components of bone, inhibiting osteoblastic differentiation and bone formation, increasing adipogenesis of MSCs at the expense of their osteogenic differentiation, and inducing osteoclastic differentiation and bone resorption (8)(40)(41). In addition, oxidized lipids induce the expression of cytokines such as *MCP-1, M-CSF* and *IL-6* both *in vitro* and *in vivo*, thought to play a role in osteoporotic bone loss (42)(43).

One of the key assumptions underlying the MR approach is that the SNPs used as genetic instrumental variables are only related to the outcome of interest through the exposure variable under study. This is invalidated by horizontal pleiotropy, whereby the genetic instrument for the exposure relates to the outcome via separate pathway to the exposure. There were several potential mechanisms for horizontal pleiotropy in the present study. For example, based on our previous finding that BMI is causally related to BMD (13), and the fact that LDL-C tends to be higher in obese individuals, it’s conceivable that relationships which we observed between LDL-C and BMD are affected by BMI. That said, whereas we observed an inverse relationship between LDL-C and BMD, increased BMI has a positive effect on BMD, suggesting these causal pathways would be in opposite directions. In addition, it’s conceivable that some of the SNPs selected from the lipid GWAS affect BMD independently of altered lipid levels. For example, a variant upstream of *ESR1* is very strongly associated with BMD (and LDL-C). Likewise a SNP in the *RSPO3* gene is strongly associated with BMD, HDL-C and triglycerides (although not with LDL-C) (44). Furthermore, a given lipid-related SNP could conceivably affect BMD by influencing a different lipid class, since many of the SNPs are associated with multiple lipid subtypes (see Figure 1).

Despite these potential sources of horizontal pleiotropy, there was not strong evidence of directional pleiotropy within the set of 147 LDL-C instruments, as reflected by the estimate of the MR-Egger intercept (I=-0.002, p=0.09), and the finding that the causal estimate from MR-Egger was consistent with estimates obtained from the IVW MR and weighted median MR methods. Moreover, we observed an almost equivalent effect of LDL-C on BMD following exclusion of obesity-associated SNPs (*β* = −0.058, 95%CI = −0.082 to −0.034, P = 3×10^-6^). In addition, there was little evidence of a causal effect of HDL-C or triglycerides on BMD, and in multivariable MR analysis the effect of LDL-C on BMD was independent of these other lipid classes. Our findings contrast with a recent MR study based on a more restricted sample of UK Biobank that found evidence of an inverse effect of BMD, as measured here, on HDL-C and other cardiovascular and type 2 diabetes mellitus risk factors (45).

A causal relationship between LDL-C and BMD is consistent with estimates of the genetic correlation between blood lipids and BMD (46) using Linkage Disequilibrium (LD) score regression (47)(48)(49). However, unlike the present analyses, a genetic correlation does not imply causality or indicate a direction of effect. Interestingly, our bidirectional MR analyses provided some evidence for a causal effect of BMD on LDL-C in all three sets of MR analyses (i.e. inverse variance, MR Egger and weighted median approaches), although large Cochrane Q statistics suggested the existence of uncontrolled genetic pleiotropy that may have contaminated the analyses. A putative causal pathway is consistent with several lines of evidence that the skeleton plays a role in regulating energy metabolism. For example, research involving mouse models has suggested that bone turnover, which is inversely related to BMD, influences insulin sensitivity and adiposity via osteocalcin (an osteoblast-specific protein) (50). Likewise, individuals with rare genetic mutations in *LRP5* which predispose to very high bone mass have markedly increased fat mass and reduced bone turnover, consistent with a causal influence of reduced osteocalcin on fat accumulation (51) Given the strong relationship between LDL-C levels and insulin sensitivity (52), our observation that BMD has a causal effect on LDL-C is consistent with this apparent control of energy metabolism by the skeleton.

### Strengths and Limitations

One of the main strengths of our study was use of two very large GWAS, which helps overcome power limitations of MR. In addition, application of two-sample MR avoids bias towards the observational association caused by weak instruments (53). A further strength of our study was the application of several analytical approaches to detect and correct for horizontal pleiotropy. One of the main limitations of the study was the use of heel ultrasound-derived BMD, in contrast to DXA-derived BMD which is used much more widely clinically to assess fracture risk. That said, heel-ultrasound derived measures have previously been found to predict fracture risk as accurately as DXA-based measures (54), and to be moderately correlated with DXA derived BMD at the hip and spine (r = 0.4 to 0.6) (55)(52).

### Conclusions

Having performed an MR study to examine the causal effect of lipid lowering on BMD, we found evidence that lowering LDL-C improves BMD, independently of HMGCR inhibition. Our results illustrate how MR can be used profitably to investigate clinical questions and drug interventions relevant to osteoporosis and bone health. Further studies are justified to explore the mechanisms by which lower LDL-C improves BMD, and to examine their potential role in treating osteoporosis, for example based on methods such as network MR (56).

## Supporting information

## Acknowledgments

NMW is supported by an NHMRC Early Career Fellowship (APP1104818). DME is supported by a National Health and Medical Research Council (NHMRC) Senior Research Fellowship (1137714). This work was supported by a Medical Research Council Programme Grant (MC_UU_12013/4 to D.M.E) and the NHMRC (1125200 to D.M.E). MAK works in a unit that is supported by the University of Bristol and UK Medical Research Council (MC_UU_12013/1). The Baker Institute is supported in part by the Victorian Government’s Operational Infrastructure Support Program. JPK is funded by a University of Queensland Development Fellowship (UQFEL1718945).

