## Supplementary material for "The effect of plasma lipids and lipid lowering interventions on bone mineral density: a Mendelian randomization study"

Supplementary Figures

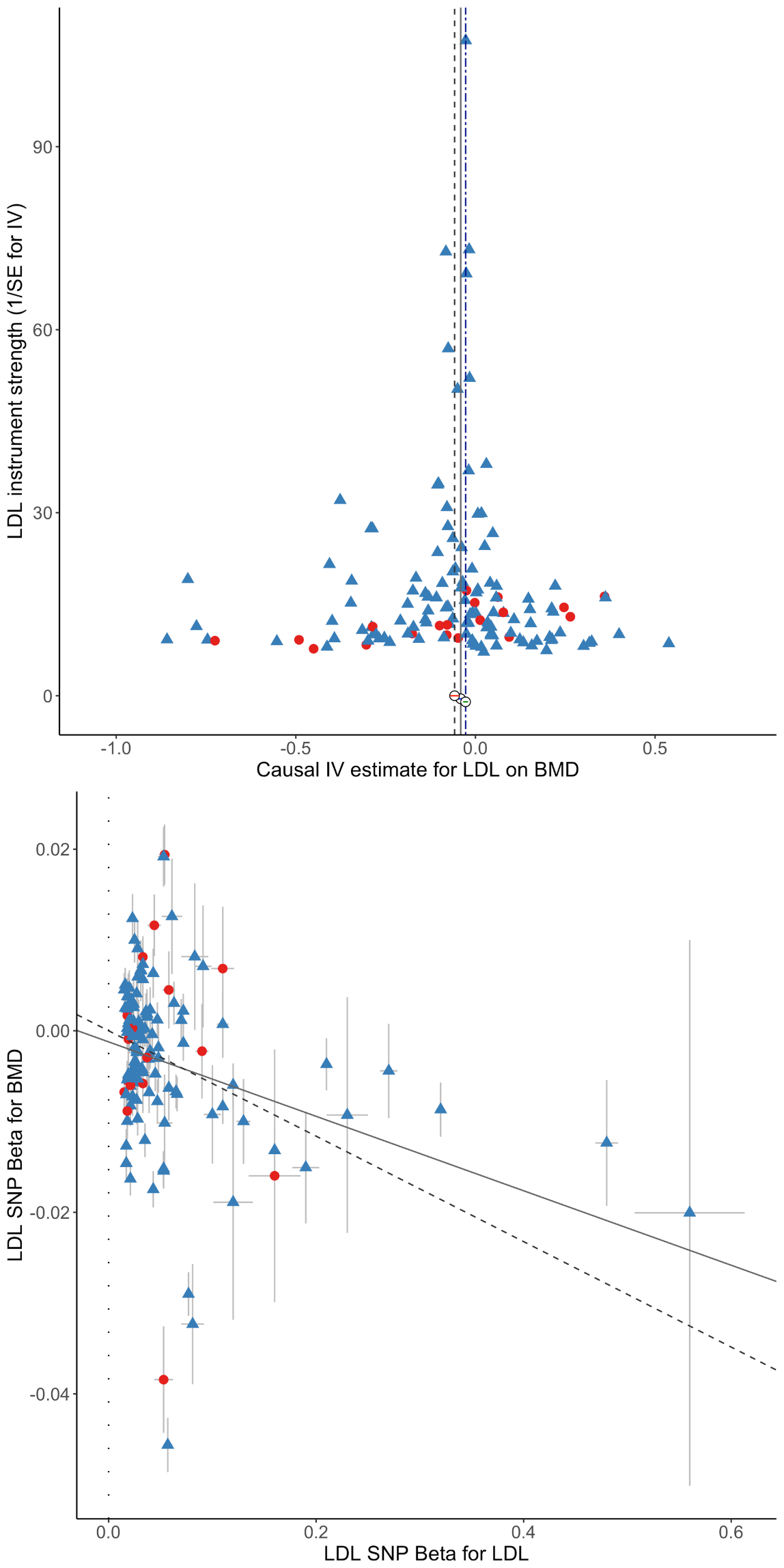

**Supplementary Figure 1.** Results for the MR analysis of LDL-C on BMD excluding BMI associated SNPs. Funnel plot (Upper plot) displaying instrument strength (y-axis) plotted against causal effect estimate (x-axis) for 137 SNPs associated with LDL-C following the exclusion of 10 BMI associated SNPs. The scatter plot (Lower plot) displays estimates of the association between each SNP and BMD (y-axis) against estimates of the association between each SNP and LDL-C level. SNPs in red circles are those associated with LDL-C (P<5x10^-8^) but not the other two lipid fractions (P>0.05), whereas the remaining SNPs are denoted by blue triangles. In the funnel plot, the inverse-variance weighted MR, MR-Egger and weighted median causal effect estimates are represented by dotted, solid and double dotted lines respectively. In the scatter plot, the slope of the solid line represents the MR-Egger regression estimate of the causal effect of LDL-C on BMD. The y-intercept of the solid regression line is an estimate of the degree of directional pleiotropy in the dataset. The inverse-variance weighted causal effect estimate is represented by the slope of the dotted line.

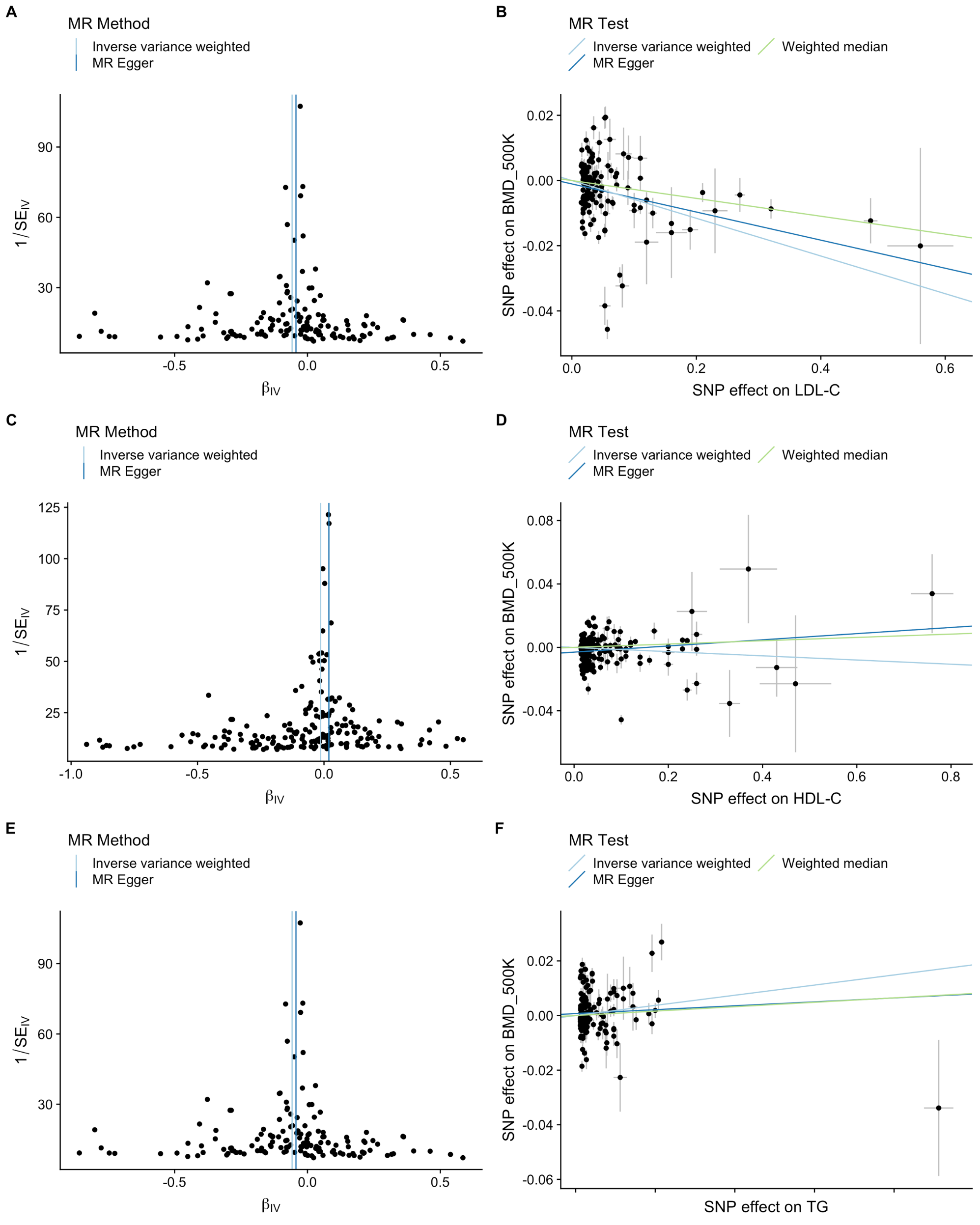

**Supplementary Figure 2.** Results of the MR analyses of lipid fractions on BMD using the Steiger filtered set of SNPs. Funnel plots on the left side of the Figure display the strength of association between each of the 146 SNPs and LDL-C (Panel A), 190 SNPs and HDL-C (Panel C) and 158 SNPs and triglycerides (Panel E) plotted against the SNP’s causal effect estimate (i.e. of the particular lipid fraction on BMD). On the right hand side of the figure are scatter plots displaying estimates of the association between each SNP and BMD against estimates of the association between each SNP and lipid fraction [i.e LDL-C (panel B), HDL-C (panel D) and triglycerides (panel F)]. The inverse variance weighted and MR Egger causal effect estimates are represented by a light blue and dark blue line respectively. The y-intercept of the dark blue regression line denotes the estimate of the degree of directional pleiotropy in the dataset. The inverse-variance weighted causal effect estimate is represented by the slope of the light blue line. The weighted median causal effect estimate is the slope of the light green line.

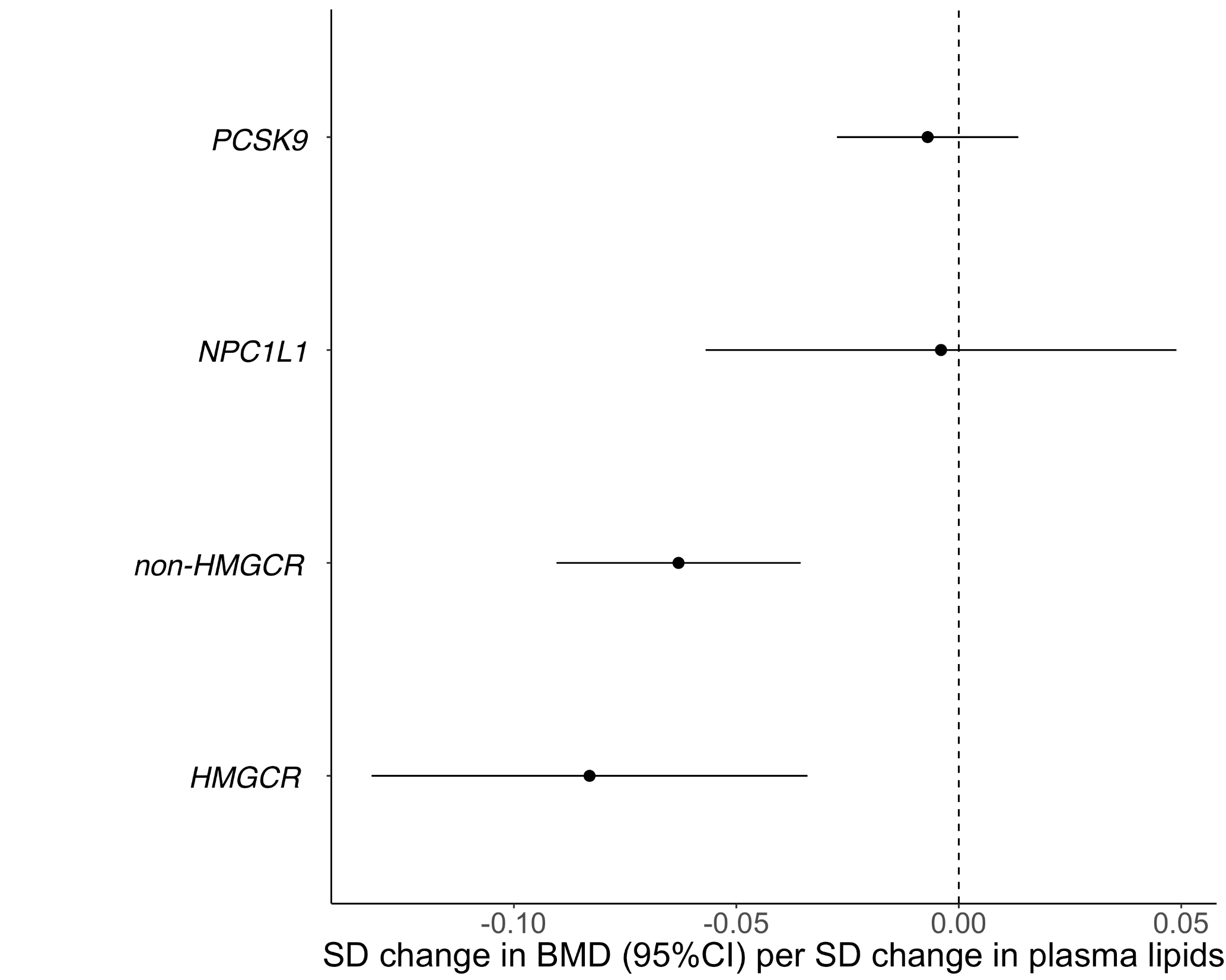

**Supplementary Figure 3.** Forest plots displaying causal estimates of the effect of LDL-C on BMD obtained using SNPs in different genomic regions [i.e. 5 *HMGCR* SNPs, 140 SNPs across the genome excluding *HMGCR*, *PCSK9* and *NPC1L1* SNPs (non-*HMGCR*), 7 *PCSK9* SNPs and 5 *NPC1L1* SNPs]. A likelihood-based approach which models the correlation between genetic variants in linkage disequilibrium was used for this analysis.

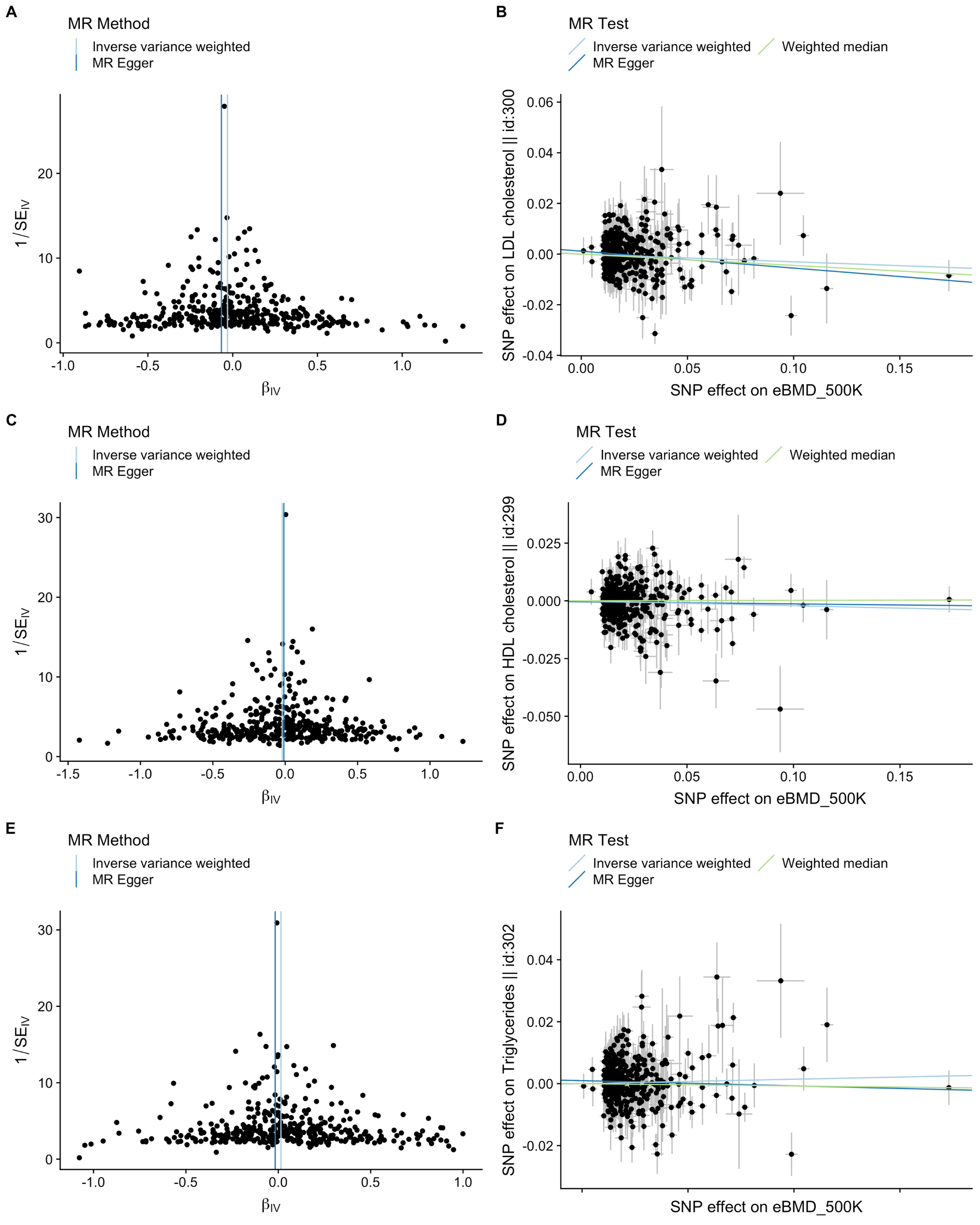

**Supplementary Figure 4.** Results of the MR analysis of BMD on lipids. Funnel plots displaying the strength of association between each of the BMD SNPs plotted against the causal estimate of the 394 SNPs on LDL-C (Panel A) and 389 SNPs on HDL-C (Panel C) and 396 SNPs on triglycerides (Panel E) and scatter plots displaying estimates of the association between each BMD SNP against effect estimates of each SNP with the relevant outcome [i.e LDL-C (panel B), HDL-C (panel D) and triglycerides (panel F)]. The inverse variance weighted and MR Egger causal effect estimates are represented by a light blue and dark blue lines respectively. The y-intercept of the dark blue regression line denotes the estimate of the degree of directional pleiotropy in the dataset. The inverse-variance weighted causal effect estimate is represented by the slope of the light blue line. The weighted median causal effect estimate is the slope of the light green line.

Supplementary Tables

Supplementary Table 1. The genetic instruments for the lipid fractions used in the multivariable Mendelian randomization analysis.

--See spreadsheet

Supplementary Table 2 to 4. List of restricted and unrestricted instruments being used for the univariate MR analyses for HDL-C, LDL-C and triglycerides representative.

--See spreadsheet

Supplementary Table 5. Estimates of causal effect of lipid fractions on BMD using multivariable Mendelian randomization

--See spreadsheet

Supplementary Table 6. MR results for SNPs in genes that are the target of lipid lowering therapies

--See spreadsheet

Supplementary Table 7. Heterogeneity test for Mendelian randomization results using SNPs within different lipid lowering target genes

--See spreadsheet

Supplementary Table 8. LD correlation matrix (r) for HMGCR, NPC1L1 and PCSK9 SNPs

--See spreadsheet

Supplementary Table 9. SNPs significantly associated with eBMD from UK Biobank and association with lipid fractions in the GLGC consortium and Steiger filtering

--See spreadsheet

Supplementary Table 10. Estimated causal effect of eBMD on lipid fractions after Steiger filtering

--See spreadsheet
